# Stainless Steel-Synthesized Gold Nanoparticles: Insights into Blood-Brain Barrier Permeability and Microglial Interaction

**DOI:** 10.1101/2025.07.30.667668

**Authors:** Zaira I. González-Sánchez, Ángela Inmaculada López-Lorente, Julia Castillo-González, Elena González-Rey, Soledad Cárdenas, David Pozo

**Affiliations:** Department of Natural and Exact Sciences, Pontificia Universidad Católica Madre y Maestra, PUCMM, Santiago de los Caballeros, Dominican Republic; Departamento de Química Analítica, Instituto Químico para la Energía y el Medioambiente (IQUEMA), Universidad de Córdoba, Campus de Rabanales, Edificio Marie Curie, E-14071, Córdoba, Spain; Department of Cell Biology and Immunology, Institute of Parasitology and Biomedicine López-Neyra (IPBLN), CSIC, Granada, Spain; Lausanne University Hospital, Lausanne, Switzerland; Department of Medical Biochemistry, Molecular Biology and Immunology, University of Seville Medical School, Av. Sanchez Pizjuan s/n, 41009, Seville, Spain; CABIMER-Andalusian Center for Molecular Biology and Regenerative Medicine (CSIC-University of Seville-UPO), Americo Vespucio s/n. Parque Científico y Tecnológico Cartuja 93, 24, 41092, Seville, Spain

**Keywords:** Gold nanoparticles, central nervous system, blood-brain barrier, microglia, interleukin-6 (IL-6)

## Abstract

Gold nanoparticles (AuNPs) are emerging as promising tools in biomedical research due to their unique physicochemical properties. However, their interactions with resident immune cells of the central nervous system (CNS) and the blood-brain barrier (BBB) remain poorly understood. This work attempts to fill this gap by evaluating the interaction of green synthetic nanoparticles -prepared using stainless steel as a sustainable reducing agent-with microglia cells, as well as their ability to permeate and potentially affect BBB properties. Using *in vitro* BBB models, we demonstrate that stainless steel-synthesized AuNPs (ss-AuNPs) effectively cross the barrier without compromising its integrity. In microglial cells, ss-AuNPs exhibited dose-dependent cytotoxic effects, with higher concentrations reducing cell viability while inducing minimal inflammatory responses. Similarly, in primary microglial cultures, ss-AuNPs at higher doses stimulated interleukin-6 (IL-6) release, although no significant pro-inflammatory activation was observed at lower doses. These results highlight the potential of green-synthesized AuNPs for CNS-targeted applications while underscoring the importance of dose optimization for safety.

## 1. Introduction

Metallic nanoparticles (NPs) have garnered significant attention in biomedical research due to their unique properties and relatively low toxicity. Gold nanoparticles (AuNPs), in particular, stand out for their chemical stability, tunable size, and possible functionalization, making them ideal candidates for applications ranging from imaging to drug delivery. However, while bulk gold is largely biocompatible, nanoscale gold exhibits distinct physicochemical and biological characteristics that require careful evaluation. Studies have shown that size, shape, surface charge, and synthesis methods significantly influence the biological interactions and toxicity of AuNPs (Ferreira et al. 2017; Jia et al. 2017; López-Lorente et al. 2019; Vijayakumar et al. 2021). For example, spherical AuNPs generally demonstrate lower cytotoxicity than spiky or rod-shaped variants, and smaller particles are often associated with increased toxicity (Hutter et al. 2010; Jia et al. 2017).

Synthesis methods also play a critical role in determining the biocompatibility of AuNPs. Conventional citrate reduction methods often result in negatively charged AuNPs, which show limited interaction with biological barriers like the blood-brain barrier (BBB) (Turkevich, et al., 1951; Kimling et al., 2006). There is growing interest in adopting greener synthesis approaches for AuNPs, due to their potential to enhance biocompatibility and reduce environmental impact (Gupta et al. 2023; Niżnik et al. 2024). Recent studies have explored innovative methods, such as the use of stainless steel as a solid reducing agent, resulting in AuNPs with different surface characteristics, particularly positive surface charge, which could lead to decreased toxicity (Han et al. 2013; López-Lorente et al. 2014a, b; Izzi et al. 2020).

Although the potential utilization of AuNPs in nanomedicine widens, there is a lack of comprehensive information related to nanoparticle-BBB interactions. The BBB, a highly selective semipermeable membrane, functions to restrict the passage of molecules and particles from the bloodstream to the CNS. There are some properties of NPs that affect this ability, such as size, surface charge, and morphology (Ruff et al. 2017; Mohammad et al. 2022; Han et al. 2025). It is known that a negative surface charge impedes particles from passing through the BBB, while making it positive promotes its transfer (Qiao et al. 2015; Ruff et al. 2017; Wu et al. 2023). NPs surface modification with peptides or proteins has been an approach to achieve BBB crossing (Prades et al. 2012; Fytianos et al. 2015; Zhang et al. 2016; Betzer et al. 2017; Ruff et al. 2017; Cox et al. 2018; Mohammad et al. 2022; Han et al. 2025), frequently changing surface charge from negative to positive.

The above-mentioned exceptional properties of metal NPs, particularly AuNPs, render them promising candidates for various biomedical applications, including cancer imaging and treatment (He et al. 2008; Hutter and Maysinger 2011; Hornos Carneiro and Barbosa 2016; Xiao et al. 2017; Kesharwani et al. 2023; Sakore et al. 2024). In particular, AuNPs can be heated by near-infrared radiation, making them valuable for hyperthermia-based treatments, particularly in brain tumor therapy (Qiao et al. 2015; Feng et al. 2017; Peng et al. 2017; Mazur et al. 2018). However, an important concern regarding the therapeutic application of AuNPs lies in their ability to cross the BBB and access the CNS. The BBB serves as a formidable defense mechanism, regulating the passage of substances between the bloodstream and the CNS to maintain brain homeostasis. Regarding NPs, size and surface charge are critical determinants of BBB penetration, with positively charged particles showing increased BBB crossing and cell internalization (Qiao et al. 2015; Ruff et al. 2017). Despite the importance of understanding interactions between metallic NPs and the BBB, available information remains limited and insufficient, particularly due to the use of oversimplified *in vitro* BBB models. To obtain physiologically relevant insights, comprehensive models should incorporate both endothelial and astrocytic cells, preferably primary astrocytes.

A significant issue for AuNPs reaching the CNS is their potential interaction with resident immune cells such as microglia, pivotal players in CNS immune defense (Battaglini et al. 2024; Gao et al. 2024). Microglia act as primary responders to pathogens and cellular damage, triggering the release of pro-inflammatory mediators and recruiting cells from the adaptive immune system, playing a dual role by initiating protective responses but also contributing to neurotoxicity (Lehnardt 2010). Microglia have the ability to sense pathogen- and host-derived ligands by engaging Toll-like receptors (TLRs), a major family of pattern recognition receptors that mediate both innate and adaptive immunity (Lehnardt 2010).

This study presents a novel approach by employing stainless steel as a sustainable reducing agent for AuNP synthesis, yielding particles with distinct surface characteristics that may enhance biocompatibility and BBB permeability. Unlike conventional citrate-based AuNPs, which exhibit limited BBB penetration due to their negative surface charge, ss-AuNPs possess a positive charge in aqueous media, potentially facilitating BBB crossing. Herein, we investigated the permeability of ss-AuNPs across two *in vitro* BBB models and evaluated their interactions with microglial cells to assess both their therapeutic potential and safety considerations. By integrating green synthesis principles with neuro-nanomedicine applications, this work provides critical insights into the development of safer and more effective nanotherapeutics for CNS disorders.

## 2. Experimental

### 2.1. AuNPs synthesis

Gold nanoparticles were synthesized mediated by stainless steel as reducing agent, following a procedure similar to that previously described (López-Lorente et al. 2019). Briefly, prior to synthesis, all the material was washed with aqua regia, *i.e*., a 1:3 v/v mixture of nitric and hydrochloric acid. Then, AuNPs were prepared by introducing an AISI 304 stainless steel disc, composed of Fe/Cr18/Ni10 with a thickness of 0.5 mm and a dimeter of 8 mm (provided by GoodFellow (Cambridge, United Kingdom) into a vial containing 200 μL of a 0.1% (w/v) solution of gold (III) chloride trihydrate (≥99.9%, Sigma Aldrich). The solution was stirred for 8 min, during which the solution’s color changed from yellow to red, confirming the formation of AuNPs.

### 2.2. AuNPs characterization

Transmission electron microscopy (TEM) was employed to visualize the generated ss-AuNPs and to calculate their average size. A JEOL JEM-1400 microscope available at the central Service for Research Support (SCAI) of the University of Córdoba (UCO) was employed. For the measurements, a drop of the ss-AuNPs solution was drop-cast on top of a copper TEM grid with a Carboward forward. ImageJ software was employed for analyzing ss-AuNPs size.

Dynamic light scattering (DLS) and Z-potential measurements were performed by using a Zetasizer-Nano ZSP (Malvern Instruments, Malvern, United Kingdom) with a 633 nm laser working at 10 mW. DLS technique was employed to calculate the hydrodynamic diameter of ss-AuNPs suspensions prepared in different media and located in a disposable polystyrene cell. The hydrodynamic diameter was calculated from the intensity distribution using a backscattering configuration with a 173º fixed-angle detector. Z-potential measurements were carried out based on laser Doppler electrophoresis and phase analysis light scattering, using a capillary cell and a forward scattering configuration.

The concentration of the ss-AuNPs was calculated by inductively coupled plasma mass spectrometry (ICP-MS), using a Perkin Elmer NexionX instrument coupled to an Argon plasma for ionization and a quadrupole detector, available at the SCAI of UCO. For ICP-MS analysis, samples were digested with 1 mL of a mixture of HNO_3_ 69% and HCl 37% 4:1 (v/v).

### 2.3. Ss-AuNPs purification

The ss-AuNPs were centrifuged at 12,000 g for 12 min and washed thrice with phosphate buffer saline (PBS). Finally, these AuNPs were resuspended to the correct concentration in PBS.

### 2.4. Cell cultures for the BBB in vitro model

Cell cultures for BBB were carried out following the methods previously described by Castillo-González et al. 2023. Murine brain endothelial cell line (bEnd.5) was purchased from Sigma. The cells were grown in b.End5 medium: Dulbecco’s Modified Eagle’s Medium (DMEM)-high glucose (4.5 g/L D-glucose) supplemented with 2 mM L-glutamine, 1 mM sodium pyruvate, 1% non-essential amino acids, 1% penicillin/streptomycin, 10% heat inactivated fetal bovine serum (FBS) (all from Gibco), and were maintained under standard cell culture conditions at 37ºC and 5% CO_2_. Medium was changed every 2 days. When sub-confluent, cells were removed from the tissue culture flask by trypsinization (with 0.25% trypsin/EDTA, Gibco) and centrifuged at 150g for 5 min at 4°C. The cell pellet was suspended in cell culture medium, and cells were maintained for successive passaging 1:4 in 75 cm^2^ flasks until used (cells between the 4^th^ to 15^th^ passage were used for experiments). This cell model has been described to be an appropriate choice to study BBB function ((Omidi et al. 2003; Yang et al. 2007)).

Primary astrocytes were obtained from P0 to P2 newborn C57BL/6 mice, as previously described (Mestre et al. 2011). Briefly, brains were dissected, and the olfactory bulb, cerebellum, hindbrain, and meninges were discarded. Then, they were homogenized in DMEM supplemented with 10% FBS, 10% horse serum (Gibco), and 1% penicillin/streptomycin. Brain homogenates were centrifuged at 1000 rpm for 10 min. The resulting cells were plated in poly-D-lysine-coated flasks (PDL, at 5 µg/mL for 30 min at 37ºC, Sigma) and incubated with astrocyte medium consisting of DMEM supplemented with 10% FBS and 1% penicillin/streptomycin for 10–12□days at 37°C and 5% CO_2_. Subsequent to microglia and oligodendrocyte progenitor cells removing, astrocytes remaining in the flasks were incubated with 12 µM cytosine β-D-arabinofuranoside (Sigma) for 3 days to avoid glial cell proliferation. Astrocytes were then trypsinized, centrifuged at 1500 rpm for 10 min, and resuspended in astrocyte medium before seeding into the transwells. Purity of astrocytes was determined by immunofluorescence as > 99% GFAP+.

### 2.5. BBB in vitro model

For both monoculture and coculture models, 24-well plate polyethylene terephthalate transwells (filter surface area 0.33 cm^2^, pore size, 8 µm; Falcon Corning Cell Culture Transwells) were coated for 60 min at 37ºC with collagen type I (50 µg/mL, dissolved in 0.02 N acetic acid; BD Falcon), washed with PBS three times and coated again for 60 min at 37ºC with fibronectin (50 µg/mL; Invitrogen). In the monoculture BBB model, b.End5 cells were seeded on the top of the filter at a density of 5 × 10^4^ cell/well in b.End5 medium. Cells were cultured at 37ºC and 5% CO_2_ until 80% confluence. In the co-culture BBB model, after extensive washing with PBS, astrocytes (5 × 10^4^ cell/well) were allowed to adhere onto the underside of the inverted filter for 15 min. Next, transwells were flipped and maintained in astrocyte medium. After 48 h (when astrocyte monolayer reached 70-80% confluence), b.End5 cells were seeded on the top of the filter as described for the monoculture model and allowed to grow for another 24 h. Contamination of adherent astrocytes on the bottom of the well was avoided by changing the transwells to another clean plate. We considered that mono and co-culture BBB models were successfully established when transendothelial electrical resistance (TEER), measured using an EVOM^2^ Epithelial Voltohmmeter (World Precision Instruments) were close to 100 Ω/cm^2^ and 200 Ω/cm^2^, respectively (Gaillard et al. 2001; Yang et al. 2007).

### 2.6. Experimental conditions for the BBB permeability assay

Once the tight barrier properties of the monolayers in the mono- and co-culture BBB models were confirmed by measuring their TEER, we added 0.1 mL of ss-AuNPs at concentrations of 1, 10, or 100 µg/mL diluted in b.End5 medium to the apical side of the BBB layers. After 24 h, the medium from both the apical and basolateral sides of the BBB models was collected. The concentration of particles at the bottom of the well was then evaluated compared to the total amount of nanoparticles added to the top. Coated transwells without cells served as positive controls. The ss-AuNPs-induced changes in the BBB permeability were analyzed by determining the flux of two permeability markers: sodium fluorescein (Na-F, MW 376 Da) and Evans blue-albumin (EBA, MW 67,000 Da), as previously described (Takata et al. 2013). Briefly, the top and bottom of the transwells were rinsed with permeability assay buffer (141 mM NaCl, 2.8 mM CaCl_2_, 1 mM MgSO_4_, 4 mM KCl, 1.0 mM NaH_2_PO_4_, 10 mM D-glucose, and 10 mM HEPES pH 7.4) and then placed in 0.6 mL of this buffer. Next, 0.1 mL of permeability buffer containing 100 µg/mL Na-F (Sigma) and 4% bovine serum albumin (Sigma) mixed with 0.67 mg/mL Evans blue dye (Sigma) was added to the apical side of the transwells, and incubated at 37ºC for 120 min. The concentration of Na-F was determined using a Tecan Infinite F200 fluorescence multiwell plate reader using a fluorescent filter pair [Ex(k) 485 ± 10 nm; Em(k) 530 ± 12.5 nm]. The EBA concentration in the abluminal chamber was measured by determining the absorbance at 630 nm with a microplate reader (Beckman, BioTek Plate Reader). The permeability coefficient (Ps, cm/min) was calculated according to the method described in (Takata et al. 2013): Ps = ([C]_A_* V_A_)/(t*S*[C]_L_), where [C]_A_ is the tracer concentration measured at the bottom chamber, V_A_ is the volume of the fluid in the bottom chamber, t is the time until tracer collection, S is the filter membrane area, and [C]_L_ is the known concentration of the tracer in the upper chamber.

### 2.7. BV-2 microglial cell culture

BV-2 microglial cell line was cultured in DMEM (high glucose) supplemented with 10% of FBS, 1% L-glutamine, and 1% penicillin/streptomycin at 37ºC in a 5% CO_2_ incubator. Cells were passaged using trypsin–ethylenediaminetetraacetic acid (EDTA) every 3–4 days.

### 2.8. Primary microglial cell culture

Primary microglial cell cultures were prepared from cerebral samples after removal of meninges of 1-to 3-day-old C57BL/6 male mice (University of Seville Animal Core Facility), and microglia were isolated by mild trypsinization as previously described (Roodveldt et al. 2013; Pérez-Cabello et al. 2023) with some modifications. After mechanical, trypsin-mediated dissociation (Lonza, BioWhittaker, Verviers, Belgium), followed by filtration in DMEM-F12 with 10% inactivated FBS (Lonza, BioWhittaker, Verviers, Belgium), cells were cultured at 37°C into 24-well plates treated with poly-D-lysine (Sigma-Aldrich, St. Louis, USA). Culture medium was carefully changed each 3 – 4 days. Cells were used for stimulation at day 18–22 of culture.

### 2.9. Cell viability assessment (MTT assay)

100 μL of BV-2 or primary microglial cells were seeded into 96-well plates at 5·10^4^ cells/mL and incubated overnight in a humidified atmosphere (37 °C, 5% CO_2_) to allow cellular adhesion. The cells were then treated with ss-AuNPs at concentrations of 1, 10, or 100 µg/mL for 24 and 48 h. After that, 10 μL of 3-(4,5-dimethylthiazol-2-yl)-2,5-diphenyltetrazolium bromide (5 mg/mL; MTT) was added to each well and incubated in the dark for 4 h. The resulting formazan crystals were then solubilized with 100 μL of dimethyl sulfoxide and incubated overnight under the same conditions. Finally, absorbance was measured using a microplate reader at 595 nm. PBS and the proteasome inhibitor MG132 were used as negative and positive controls, respectively.

### 2.10. Lactate dehydrogenase (LDH) release assay

Extracellular LDH release and total LDH were evaluated in cells treated as described above. After treatment, the supernatants of lysed (total LDH) or not lysed cells (extracellular LDH) were used in the determination with Cytotoxicity Detection Kit Plus (LDH, Roche) following the manufacturer’s instructions. In brief, 100 μL of reaction mixture was added to each well and incubated for up to 30 min. Then, 50 μL of stop solution was added, and the absorbance was measured at 490 nm in a microplate reader.

### 2.11. TLR stimulation

For TLR stimulation, 500 μL of BV-2 cells were seeded in 24-well plates at a density of 9·10^5^ cells/mL and incubated overnight in a humidified atmosphere (37°C, 5% CO_2_) to allow cellular adhesion. Cells were treated for 24 h with the specific TLR ligand at optimal concentration in the presence or absence of 1, 10, or 100 µg/mL of ss-AuNPs. TLR ligand final concentrations were as follows: lipopolysaccharide (LPS; 1 μg/mL, *Escherichia coli* serotype 0127:B8; Sigma, St. Louis, MO, USA) and synthetic bacterial lipopeptide Pam3Csk4 (300 ng/mL; InvivoGen, San Diego, CA, USA).

### 2.12. Microglia IL-6 secretion

After 24 h of TLR stimulation, supernatants from BV-2 cells were harvested, and IL-6 secretion was determined by enzyme-linked immunosorbent assay (ELISA) according to the manufacturer’s instructions (OptEIA Mouse IL-6 set, BD Pharmingen, San Diego, CA, USA).

### 2.13. Cell internalization assay by immunofluorescence

BV-2 or primary microglial cells were seeded in 12-well plates at 5·10^4^ cells/mL and incubated overnight at 37ºC with 5% CO_2_ to allow cell adhesion. Then, cells were treated with 100 µg/mL ss-AuNPs for 24 h. Cells were trypsinized, washed three times, and resuspended in complete medium DMEM high glucose. Cells were then seeded onto 19 mm coverslips treated with poly-D-lysine and incubated for 24 h to allow cell adhesion. After that period, coverslips were rinsed three times with PBS and samples were fixed with 4% paraformaldehyde (PFA) for 15 min at 4ºC and washed three times with PBS. Then, samples were blocked with 3% bovine serum albumin (BSA) and 0.5% Triton X-100 in PBS for 1 h at room temperature. Next, samples were incubated with Phalloidin Alexa Fluor 488 (Sigma) for 20 min at room temperature and washed three times with PBS. Finally, 4′,6-diamidino-2-phenylindole (DAPI) (Sigma) was added and incubated for 5 min at room temperature, and samples were washed three times with PBS and mounted on microscope slides using Vectashield mounting medium (Vector Laboratories, Newark, CA, USA) for fluorescence.

### 2.14. Statistical evaluation

All experiments were conducted in triplicate, including the appropriate controls. Data are presented as mean +/-standard deviation (SD). Statistics were calculated using a Student’s t-test or ANOVA and Tukey’s post-hoc test. Significance was accepted at P<0.05.

## 3. Results and discussion

### 3.1. Synthesis and characterization of ss-AuNPs

Green chemistry is coming of age, with growing interest in both academic and industrial laboratories. Green synthesis of nanomaterials aims to eliminate hazards at the design stage from the outset to create better safer chemicals while choosing the most secure and efficient ways to synthesize them and to reduce wastes. In the present work, AuNPs were synthesized using stainless steel as solid reducing agent, which leads to a different environment surrounding the AuNPs as compared with the classical reduction with citrate (López-Lorente et al., 2019). The ss-AuNPs prepared through this green method were characterized using different techniques, including TEM, DLS and Z-potential, in order to calculate their size and size distribution, as well as surface properties.

As can be observed in the TEM photographs (Figure 1A) of the synthesized AuNPs, they possess a nearly spherical shape with an average diameter of 37±17 nm, displaying a Gaussian size distribution as shown in Figure 1B. The hydrodynamic size of the ss-AuNPs dispersed in the different media (*i.e*., water, PBS and DMEM culture medium) was calculated by DLS. Figure 1B shows the average size distribution by intensity of ss-AuNPs diluted in Milli-Q water and of ss-AuNPs after purification by centrifugation and resuspended in DMEM and PBS media. The average size (by intensity) of the ss-AuNPs in the different media was 130±2, 133.0±0.7, and 719±15 nm for aqueous solution, DMEM and PBS, respectively (Figure 1C). This discrepancy likely arises from aggregation and protein corona formation in biological media, a phenomenon that can influence nanoparticle stability and bioavailability. In addition, Z-potential measurements were also performed to calculate the surface charge of the ss-AuNPs in the different media. As expected, ss-AuNPs in aqueous diluted media reveal positive charge, in agreement with previous works (Han et al. 2013; López-Lorente et al. 2019; Izzi et al. 2020). ss-AuNPs evidence highly positive Z-potential values above 30 mV, while after dilution with water, it is reduced to 16.1±0.7 mV. On the other hand, once the ss-AuNPs are centrifuged and resuspended in DMEM and PBS media, the Z-potential turned negative with values of -11.6±0.3 and -15.7±0.7, respectively. This shift suggests significant adsorption of serum proteins, which may alter cellular interactions and uptake.

**Figure 1.**
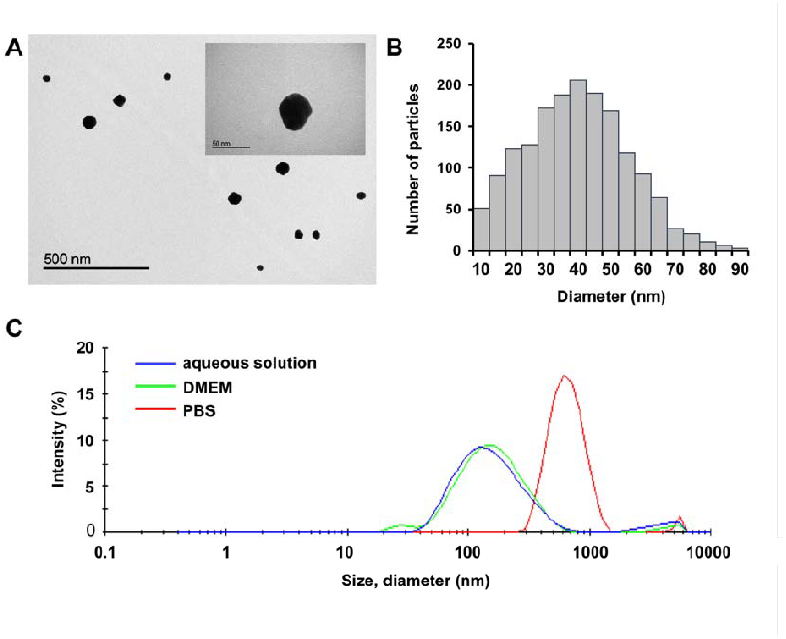
Characterization of the AuNPs synthesized via stainless steel. (A) TEM images of the individual ss-AuNPs. The inset shows one AuNP at higher magnification. (B) Size distribution of the ss-AuNPs obtained from the TEM images; (C) DLS size intensity distribution by intensity of the ss-AuNPs in different media, namely as-synthesized aqueous diluted solution (blue), DMEM (green) and PBS media (red).

### 3.2. ss-AuNPs cross the BBB in in vitro models

Although the potential applications of AuNPs in nanomedicine continue to expand, there is a lack of comprehensive information on their interactions with the BBB. The BBB restricts the passage of molecules and particles from the bloodstream to the central nervous system (CNS), posing a significant challenge for drug delivery to the brain. While many nanoparticles have been shown to cross the BBB, physicochemical properties such as size, surface charge and morphology (Ruff et al. 2017; Mohammad et al. 2022; Han et al. 2025) critically influence their ability to cross this barrier. For instance, a negative surface charge generally hinders particles from passing through the BBB, whereas a positive charge promotes transfer (Qiao et al. 2015; Ruff et al. 2017; Wu et al. 2023). Indeed, NPs surface modification with peptides or proteins has been a strategy to achieve BBB crossing (Prades et al. 2012; Fytianos et al. 2015; Zhang et al. 2016; Betzer et al. 2017; Ruff et al. 2017; Cox et al. 2018; Mohammad et al. 2022; Han et al. 2025), often shifting the surface charge from negative to positive. The AuNPs synthesized through stainless steel used in this work exhibit a positive charge in aqueous medium, as demonstrated by the Z potential measurements, but the charge becomes negative in DMEM culture medium, possibly due to a result of the formation of a protein corona.

Thus, to evaluate the ability of these ss-AuNPs to cross the BBB, two *in vitro* models were used. Both were assembled on 24-well plates with a transwell that divided each well in top and bottom compartments. The first model consisted of a monoculture of mouse endothelial bEnd.5 cultured as a confluent monolayer on the apical side of the transwell (Figure 2A, top). The second model consisted of a more physiologically relevant coculture, including primary astrocytes seeded on the underside of the inverted transwell (Figure 2A bottom). Ss-AuNPs were applied on the top side and incubated for 24 h. Subsequently, the Au concentration in the bottom compartment of each well was determined by ICP-MS, representing the quantity of ss-AuNPs that traversed the barrier. As expected, significantly more ss-AuNPs traversed the barrier in the monoculture model lacking astrocytes across all tested concentrations (Figure 2B, black bars).

**Figure 2.**
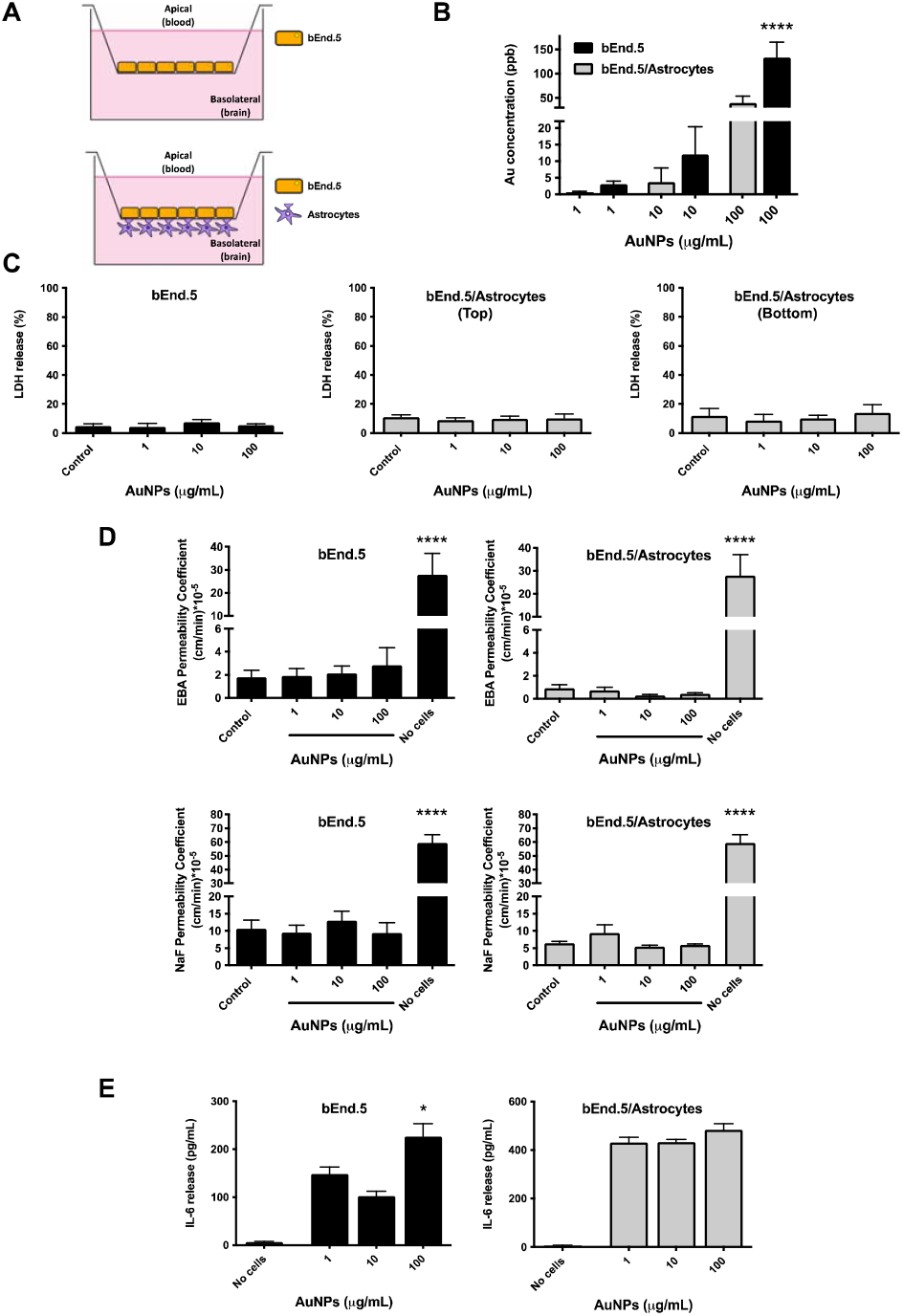
ss-AuNPs cross the BBB *in vitro*. (A) Illustration of the two *in vitro* models of BBB used in this work. (B) Au concentration on the bottom side of the barrier, which represents the ss-AuNPs that crossed the barrier. (C) Percentage of the LDH release after the treatment with different concentrations of ss-AuNPs. (D) BBB integrity after the treatment with the ss-AuNPs measured by Evans blue albumin (EBA, Top) and sodium fluorescein (NaF, Bottom) permeability. (E) IL-6 release of bEnd.5 endothelial cells (left) or bEnd.5 and astrocytes (right).

Consistent with previous findings (Ruff et al. 2017; Enea et al. 2019; Gonzalez-Carter et al. 2019; Duan et al. 2024), ss-AuNPs have the ability to cross the BBB in *in vitro* models (Figure 2B), potentially due to the impact of AuNPs on endothelial cell tight-junction integrity, as previously reported (Li et al. 2015). To ascertain whether ss-AuNPs compromise cell viability, extracellular LDH release was assessed in the supernatants of each well. No discernible differences in LDH release were observed between ss-AuNPs-treated and untreated transwells (Figure 2C), indicating an absence of cytotoxicity, consistent with findings for similar AuNPs in previous studies (Lee et al. 2019; Sobska et al. 2021). Furthermore, the integrity of the BBB was evaluated by Evans blue albumin (EBA) and sodium fluorescein (NaF) assays. The top supernatants were replaced with EBA or NaF, and fluorescence intensity was measured on the bottom side after 120 min. No significant differences were detected between ss-AuNPs-treated or untreated transwells (Figure 2D), indicating that ss-AuNPs did not compromise the integrity of the barrier.

Additionally, to assess whether the presence of ss-AuNPs prompts an immune response in endothelial cells, astrocytes, or both, IL-6 levels were measured in the supernatants. As shown in Figure 2E, treatment with 100 μg/mL ss-AuNPs increased IL-6 release in endothelial cell culture. However, no significant differences were observed in the co-culture.

BBB permeability assays in two distinct *in vitro* models confirmed that AuNPs effectively cross the barrier without compromising its integrity. These findings are consistent with previous studies highlighting the role of positively charged nanoparticles in facilitating BBB crossing (Ruff et al. 2017; Enea et al. 2019). However, unlike many traditional AuNP synthesis methods that employ surface modifications to enhance BBB penetration (Fytianos et al. 2015; Zhang et al. 2016), our ss-AuNPs achieve efficient penetration inherently, showcasing a unique biophysical interaction profile. This represents a significant advancement in the design of nanomaterials with intrinsic BBB-translocation abilities.

### 3.3. Ss-AuNPs cytotoxicity and internalization in microglial cells

Upon confirming the ability of ss-AuNPs to cross the BBB *in vitro*, we sought to evaluate their impact on microglia. To investigate how microglial cells respond to the presence of ss-AuNPs, two models were used: the BV-2 microglial cell line and primary microglial cultures from newborn mice. Both cell types were exposed to increasing concentrations of ss-AuNPs (*i.e*., 1, 10, 100 μg/mL), and cell viability, LDH release, and IL-6 secretion were assessed.

Initially, an immunofluorescence assay was conducted to evaluate ss-AuNPs internalization, using a confocal microscope. The results revealed ss-AuNPs internalization by BV-2 cells following 24 h treatment of ss-AuNPs (Figure 3A). Phalloidin binds to the α-actin of the plasma membrane, marking the contour of the cell in each plane, thus allowing the discrimination of the components inside and outside the cell. DAPI marks the nucleus (in blue), while the ss-AuNPs exhibit autofluorescence that is seen in red (Figure 3A, left images). Finally, a maximum intensity Z-projection was constructed to represent the ss-AuNPs accumulated in the different z planes (Figure 3A, right image). This internalization phenomenon of AuNPs has been previously demonstrated in various cell types, including microglia (Hutter et al. 2010; Stojiljković et al. 2016) and macrophages (Krpetić et al. 2010; Chen and Gao 2017), suggesting a potential role in cellular protective and cleaning functions.

**Figure 3.**
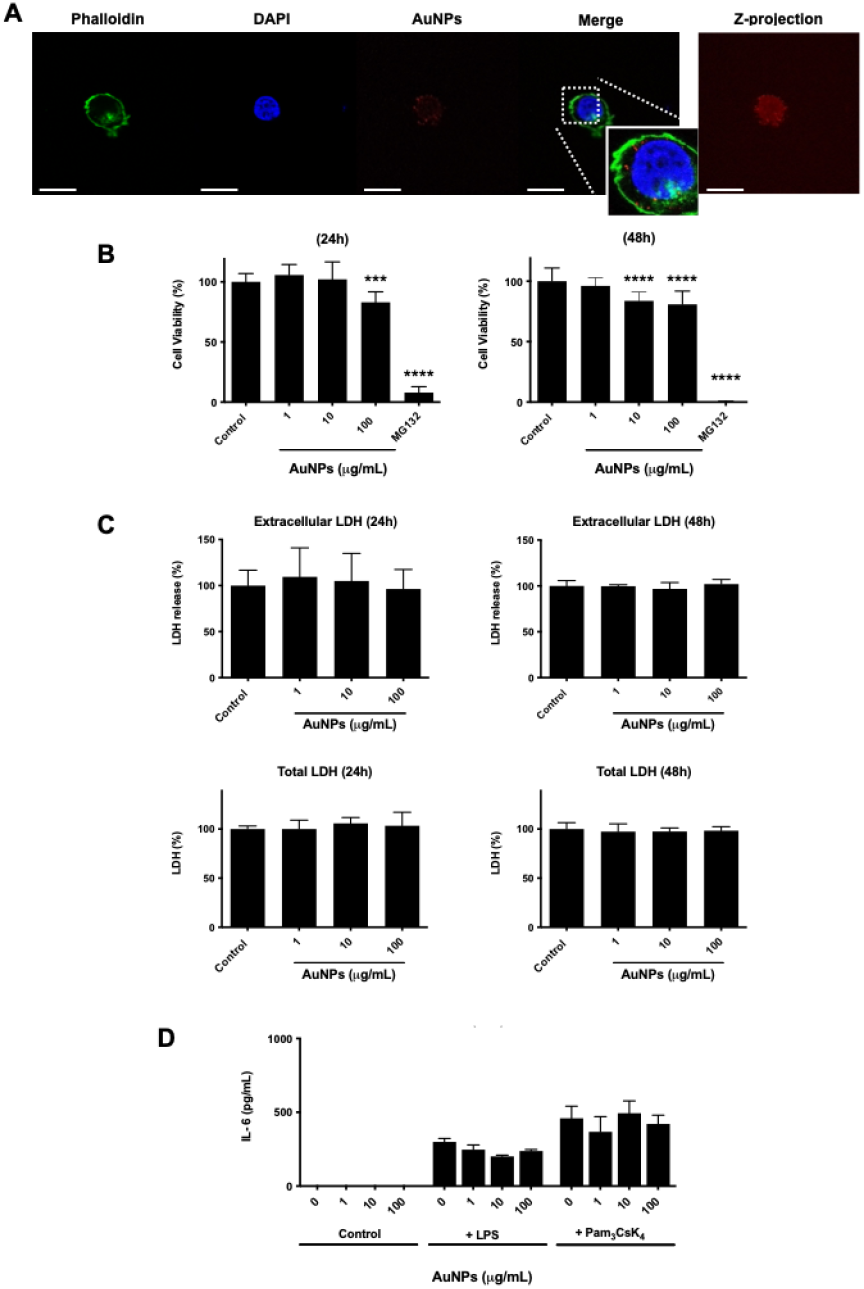
ss-AuNPs are internalized and affect the viability of BV-2 microglial cells at the highest concentration. (A) ss-AuNPs internalization observed by immunofluorescence in a confocal microscope. Phalloidin (green) was used to stain the plasma membrane and DAPI (blue) to stain the nucleus. Autofluorescence of ss-AuNPs is observed in red. Scale bar: 10 µm. (B) BV-2 microglial cell viability in the presence or absence of ss-AuNPs. MTT assay was conducted after 24 h (left) or 48 h (right) after treatment with 1, 10, or 100 µg/mL of AuNPs. PBS and MG132 were used as negative and positive controls, respectively. Values are the mean ± SD of the different replicates. (C) Extracellular and total LDH levels for BV-2 microglial cells treated with ss-AuNPs. Cells were incubated in the presence of 1, 10, or 100 µg/mL of ss-AuNPs for 24 h (top left, top right) or 48 (bottom left, bottom right) before the LDH determination. PBS was used as a negative control. Values are the mean ± SD of the different replicates. (D) IL-6 secretion in the presence of ss-AuNPs in activated or not activated BV-2 microglial cells. Cells were stimulated by LPS or CsK4. Values are the mean ± SD of the different replicates.

Assessment of the cytotoxicity of ss-AuNPs in BV-2 microglial cells consisted of measuring cell viability and LDH release. The MTT assay (Figure 3B) indicated that BV-2 cell viability was not affected by ss-AuNPs at 1 and 10 μg/mL in the first 24 h, however after 48 h of treatment both 10 and 100 μg/mL significantly affected their cell viability. These results are consistent with previous research in microglial (Hutter et al. 2010; Xue et al. 2019) and RAW 267.4 macrophage cells (Singh et al. 2017; Ahn et al. 2018). Particularly, Hutter et al., 2010 evaluated the cell viability of N9 microglial cells and primary cell culture after the treatment with gold nanospheres, nanorods and nanourchins, showing significant differences from 10^9^ NPs/mL. Similarly, Xue et al., assessed cell viability of AuNPs biosynthesized from the rhizome of *Paeonia moutan* (PM-AuNPs) on BV-2 cells, observing a cytotoxic effect starting at 10 μg/mL incubated for 24 h.

Furthermore, ss-AuNPs at these concentrations showed no effect on extracellular LDH release in BV-2 treated at 24 h or 48 h (Figure 3C, top). Similarly, no significant changes in total LDH were observed for BV-2 cells treated with the ss-AuNPs at 24 and 48 h (Figure 3C, bottom), which also agrees with recent reports (Jia et al. 2017; Cox et al. 2018; Sobska et al. 2021). Overall, our results suggest that ss-AuNPs at 1 μg/mL do not affect microglia cell viability and integrity and ensure that the changes observed below are not due to cell number.

Exploration of interleukin secretion as a marker of immune cell inflammatory response revealed that ss-AuNPs did not induce IL-6 release in either non-activated or activated BV-2 microglial cells (Figure 3D). This lack of IL-6 release suggests the absence of microglial cell activation, a finding further supported by the maintenance of cellular morphology following ss-AuNPs treatment (data not shown). Previous studies have reported similar outcomes, indicating that certain configurations of AuNPs, such as sphere-shaped ones, do not significantly activate microglia or induce cytokine release (Hutter et al. 2010; Chen and Gao 2017). Conversely, AuNPs synthesized by using plant extracts appear to exhibit anti-inflammatory activity by reducing IL-6 and other interleukins (Park et al. 2019; Xue et al. 2019). Overall, these results, combined with the observed internalization of ss-AuNPs, suggest that BV-2 microglial cells effectively clear ss-AuNPs as part of their debris-cleaning function, impacting on cell viability only at high concentrations unlikely to be reached *in vivo*.

To ensure the reproducibility of our findings in a more physiologically relevant model, primary microglial cell cultures were employed. Subsequently, the internalization of ss-AuNPs was assessed via immunofluorescence, with images captured using a confocal microscope. As depicted in Figure 4, primary microglial cells were found to have internalized ss-AuNPs following 24 h treatment with 100 µg/mL of ss-AuNPs.

**Figure 4.**
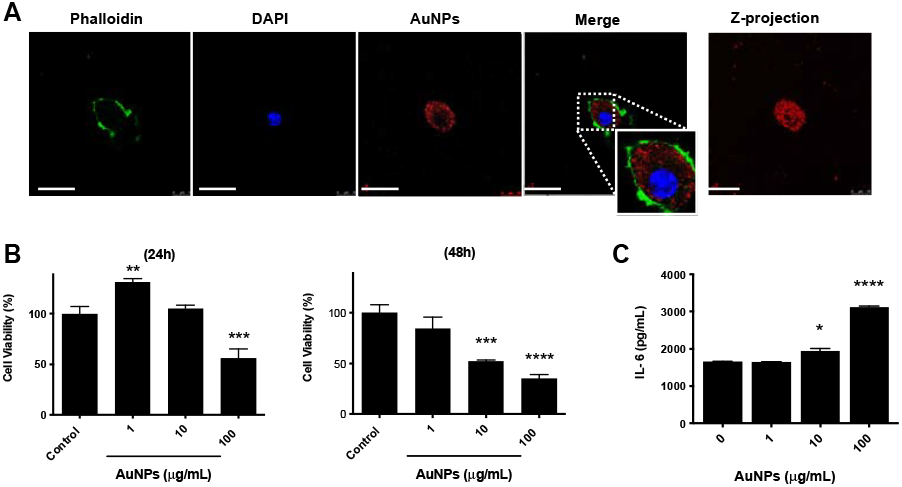
ss-AuNPs are internalized by primary microglial cells and affect their viability and cytokine release. (A) ss-AuNPs internalization in microglia primary cells. Phalloidin (green) was used to stain the plasma membrane and DAPI (blue) to stain the DNA. Autofluorescence of ss-AuNPs is observed in red. Scale bar: 10 µm. (B) Cell viability of primary microglia after treatment with different concentrations of ss-AuNPs for 24 and 48 h. MTT assays were conducted after 24 h (left) or 48 h (right) of the treatment with 1, 10, or 100 µg/mL of ss-AuNPs. Values are the mean ± SD of the different replicates. (C) IL-6 release in primary cell microglia cultures after treatment with 1, 10, or 100 μg/mL of ss-AuNPs for 24h. Values are the mean ± SD of the different replicates (N=3).

Then, primary microglial cell cultures were treated with different concentrations of ss-AuNPs and cell viability was determined by MTT. As illustrated in Figure 4B, the AuNPs synthesized via stainless steel exerted a dose-dependent effect on primary microglial cell viability, consistent with prior findings in Raw 264.7 macrophages, where 100 µg/mL of 10-30 nm AuNPs synthesized using an aqueous extract of *Angelicae Pubescentis* had an impact on cell viability (Markus et al. 2017).

In addition, our study sought to explore whether ss-AuNPs have an impact on IL-6 release in primary microglia. Treatment with ss-AuNPs at concentrations of 10 and 100 µg/mL significantly increased IL-6 release after 24 h, as depicted in Figure 4C. This observation aligns with *in vivo* findings reported by various researchers. For example, citrate-capped 20 nm Au-NPs were found to elevate the production of anti-inflammatory cytokines (IL-10; IL-4) (Liu et al. 2013), while 5 nm AuNPs were observed to decrease IL-1β on day one and enhance IL-1β and IL-6 seven days post-treatment (Khan et al. 2018). Additionally, (Khan et al. 2018) reported that a single injection of 5 nm AuNPs significantly increased mRNA expressions of IL-1β and IL-6 in the liver, with normalization observed on day 7. Moreover, in the spleen, 5, 20, and 50 nm GNPs were found to significantly increase IL-1β and IL-6 mRNA expressions on day 1, persisting through day 7 (Khan et al. 2018).

The interactions of ss-AuNPs with microglia further revealed critical dose-dependent effects on cell viability and cytokine release. Importantly, ss-AuNPs at lower concentrations induced no significant changes in BV-2 microglial cells or primary microglial cultures, supporting their biocompatibility. However, higher concentrations compromised cell viability, aligning with studies that describe size- and dose-dependent toxicity of AuNPs in immune and neuronal cells (Jia et al., 2017; Xue et al., 2019). Interestingly, unlike other studies where AuNPs provoked significant inflammatory responses via cytokine upregulation (Khan et al., 2018), our results showed minimal IL-6 release except at the highest concentration in primary microglia. This suggests that the stainless-steel synthesis method may yield nanoparticles with reduced pro-inflammatory potential, making them promising candidates for therapeutic use in the CNS.

Nevertheless, our findings underscore the importance of careful dosage consideration when applying ss-AuNPs in biomedical settings. Whereas their ability to reach the CNS without impairing BBB function is advantageous, their cytotoxicity at higher concentrations and potential to stimulate inflammatory pathways warrant further investigation. Comparatively, our results highlight the dual-edged nature of nanomedicine: the therapeutic promise of ss-AuNPs must be weighed against potential safety concerns.

## 4. Conclusion

This study highlights the promising potential of ss-AuNPs synthesized via a sustainable, steel-mediated method for crossing the BBB and interacting with microglial cells. This one-pot method used to produce the ss-AuNPs is simple, fast, and cost-effective, yielding stable AuNPs positively charged, without the presence of ligands at their surface. The ss-AuNPs demonstrated efficient permeability across the *in vitro* BBB model and were well-tolerated by both immortalized and primary microglia at low concentrations, with minimal cytotoxicity and inflammatory responses. However, higher doses induced significant immune activation, underscoring the critical importance of dose optimization. These findings suggest that green-synthesized ss-AuNPs could serve as valuable tools for neurotherapeutic delivery if carefully tuned to avoid adverse microglial reactivity. Future research should focus on *in vivo* validation of these effects and the design of controlled-release systems to harness their full therapeutic potential in CNS disorders.

## Acknowledgements

Z.I. González-Sánchez wishes to thank Ministry of Science and Technology, Dominican Republic, for funding Projects FONDOCyT 2014-1B3-051 and 2015-3A6-138 to Z.I.G-S. Z.I. This work was also supported by the Spanish Ministry of Science and Innovation (MCIN)/AEI/ 10.13039/501100011033 and by “ERDF A way of making Europe” grants PID2020–119638RB-I00 (to E.G-R.) and by FPU-program FPU17/02616 (to J.C-G.)

## Data statement

The data presented in this article would be available under request.

## Green-Synthesized Gold Nanoparticles: BBB Permeability and Microglial Response

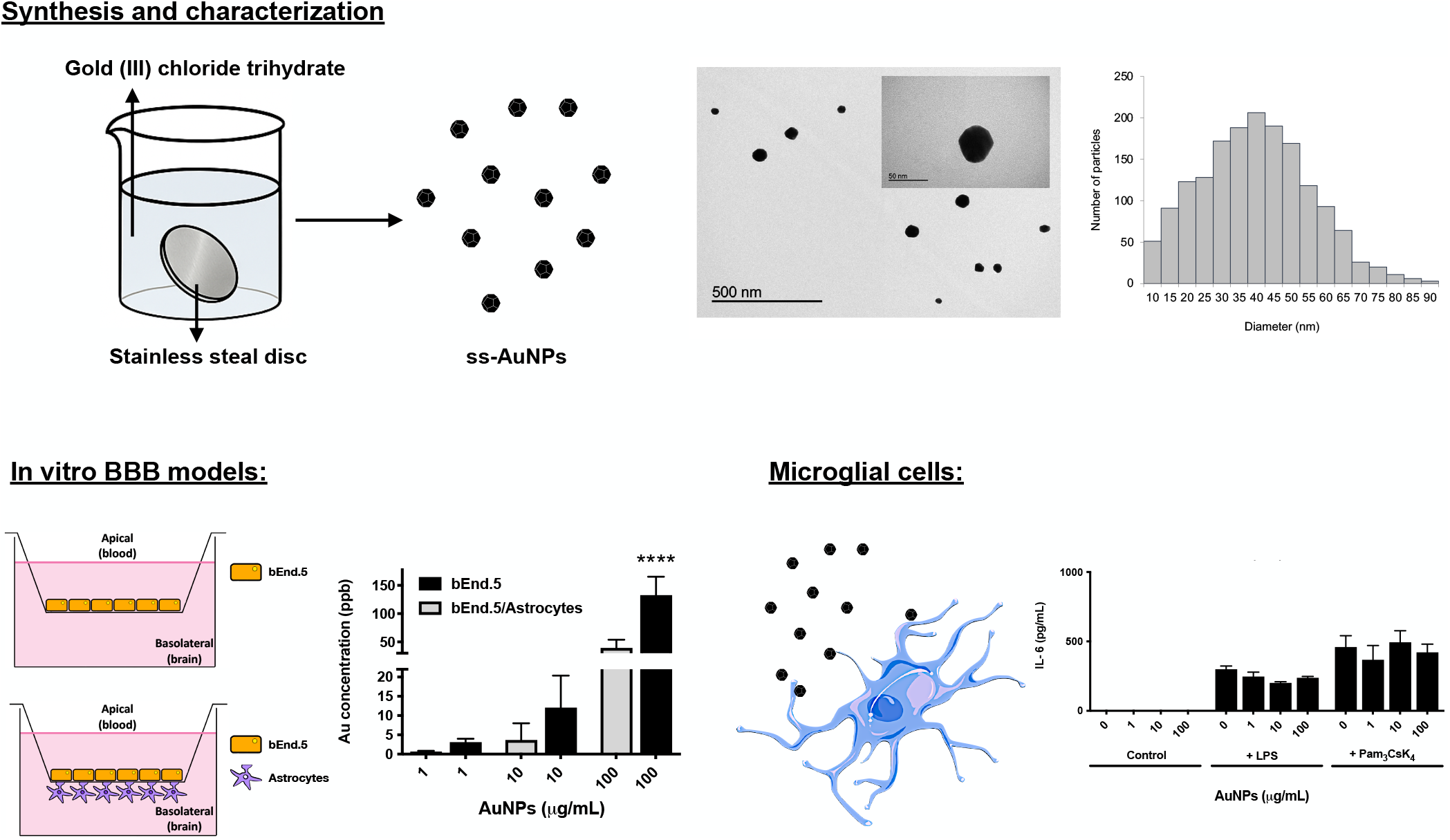

